# Identifying mixed *Mycobacterium tuberculosis* infection and laboratory cross-contamination during Mycobacterial sequencing programs

**DOI:** 10.1101/344853

**Authors:** David H Wyllie, Esther Robinson, Tim Peto, Derrick W Crook, Adebisi Ajileye, Priti Rathod, Rosemarie Allen, Lisa Jarrett, E Grace Smith, A Sarah Walker

**Affiliations:** Nuffield Department of Medicine, John Radcliffe Hospital, Headley Way, Oxford OX3 9DU, UK; Public Health England Academic Collaborating Centre, John Radcliffe Hospital, Headley Way, Oxford OX3 9DU, UK; The National Institute for Health Research Health Protection Research Unit (NIHR HPRU) in Healthcare Associated Infections and Antimicrobial Resistance at University of Oxford; Public Health England National Mycobacteriology Laboratory North and Central, Heartlands Hospital, Birmingham B9 5SS

**Keywords:** *Mycobacterium tuberculosis*, next generation sequencing, quality control, mixed infection, cross-contamination

## Abstract

**Introduction:** Detecting laboratory cross-contamination and mixed tuberculosis infection are important goals of clinical Mycobacteriology laboratories.

**Objectives:** To develop a method detecting mixtures of different *M. tuberculosis* lineages in laboratories performing Mycobacterial next generation sequencing (NGS).

**Setting:** Public Health England National Mycobacteriology Laboratory Birmingham, which performs Illumina sequencing on DNA extracted from positive Mycobacterial Growth Indicator tubes.

**Methods:** We analysed 4,156 samples yielding *M. tuberculosis* from 663 MiSeq runs, obtained during development and production use of a diagnostic process using NGS. Counts of the most common (major) variant, and all other variants (non-major variants) were determined from reads mapping to positions defining *M. tuberculosis* lineages. Expected variation was estimated during process development.

**Results:** For each sample we determined the non-major variant proportions at 55 sets of lineage defining positions. The non-major variant proportion in the two most mixed lineage defining sets (F2 metric) was compared with that in the 47 least mixed lineage defining sets (F47 metric). Three patterns were observed: (i) not mixed by either metric, (ii) high F47 metric suggesting mixtures of multiple lineages, and (iii) samples compatible with mixtures of two lineages, detected by differential F2 metric elevation relative to F47. Pattern (ii) was observed in batches, with similar patterns in the H37Rv control present in each run, and is likely to reflect cross-contamination. During production, the proportions of samples in each pattern were 97%, 2.8%, and 0.001%, respectively.

**Conclusion:** The F2 and F47 metrics described could be used for laboratory process control in laboratories sequencing *M. tuberculosis*.

## INTRODUCTION

*Mycobacterium tuberculosis* is an organism which has co-evolved with humans during the early migrations of modern man, diverging from a common *M. tuberculosis* ancestor about 75,000 years ago (1). Distinct lineages, corresponding to evolution occurring during these early migrations, are readily identified by next generation sequencing (NGS), with each lineage characterised by ancient single nucleotide variants (SNV) which define deep branches in the *M. tuberculosis* phylogeny (1, 2).

### Multiple *M. tuberculosis* lineages

Infection by multiple lineages of TB is well described, and has been detected by observing mixed results on fractional sequencing (e.g. spoligotyping, MIRU-VNTR) and validated by characterisation of multiple individual picks from solid media (3). Multi-lineage infection is characterised by isolates differing by many hundreds of SNVs, in which respect it differs from the increasingly recognised and more common in-host microevolution (4). Reported rates of mixed infection vary markedly as reviewed (3, 5), with between 10 and 30% reported in areas of current (5–7) or historical (8, 9) high prevalence. Much lower rates are reported in low incidence countries (3), although systematic under-detection is likely to occur due to both to the limited representation of bacteria in single sputum samples of pulmonary disease, and to the decrease in diversity occurring during differential strain growth in broth culture (10).

### Implications of mixed infection

Mixed infection, assessed by either MIRU-VNTR polymorphisms (7) or by heterogeneity in drug susceptibility testing (11), is independently associated with reduced drug treatment response, so there are compelling clinical reasons to try to identify it. There are also important technical implications of isolating mixed TB strains from a culture. Firstly, such a finding may reflect cross-contamination within the laboratory (3). Secondly, mixed infection complicates the interpretation of drug resistance tests, whether phenotypic or genotypic, as one or other co-infecting strain may dominate the results from these tests. Thirdly, it complicates the understanding of relatedness when techniques such as SNV distance computation are applied, as these generally assume a single sequence is present when base-calling (5, 12–14), marking mixed sites as uncertain. Maximal likelihood tree drawing algorithms assume such ‘uncertain’ sites contain no information and impute a single nucleotide at each of such positions, an approach which may be inappropriate in the presence of mixtures.

Increasingly, NGS based species and resistance determination is becoming routine in Mycobacteriology laboratories, and has been deployed in reference laboratories in England (15, 16). As part of the quality control and accreditation of the routine process now operating in these laboratories, we describe an approach to identifying mixed samples using Illumina next generation sequencing data, illustrating its use by studying over 4,000 consecutive positive cultures from a single reference laboratory.

## METHODS

### Isolation of DNA from Mycobacteria and sequencing

This study includes all mycobacteria processed between 01/06/2015 and 30/12/2017 in the Public Health England Midlands and North reference laboratory, whose catchment is approximately 15 million people, or about one third of England. Clinical specimens were decontaminated and inoculated into Mycobacterial Growth Indicator Tubes (MGIT) tubes. During the process (Fig. 1) positive MGIT tubes were batched when they became available, either following growth in the local laboratory, or following receipt from another laboratory. Positive control samples (H37Rv strain) were also grown in MGIT cultures. Batches of positive samples were extracted using a manual process, exactly as described in the Supplementary Methods in (16). Illumina sequencing libraries were prepared using Nextera XT chemistry from equal amounts of 12 (20.04.15-01.08.15) or 16 (01.08.15-30.12.17) Mycobacterial DNA extracts (16), using manual steps (Table 1). Positive control DNA (H37Rv, obtained from ATCC) was included as one of the 12 or 16 extracts in all libraries, either from a contemporaneously extracted broth culture, or from stored DNA. Libraries were loaded into an Illumina MiSeq instrument and sequenced (16).

**Figure 1.**
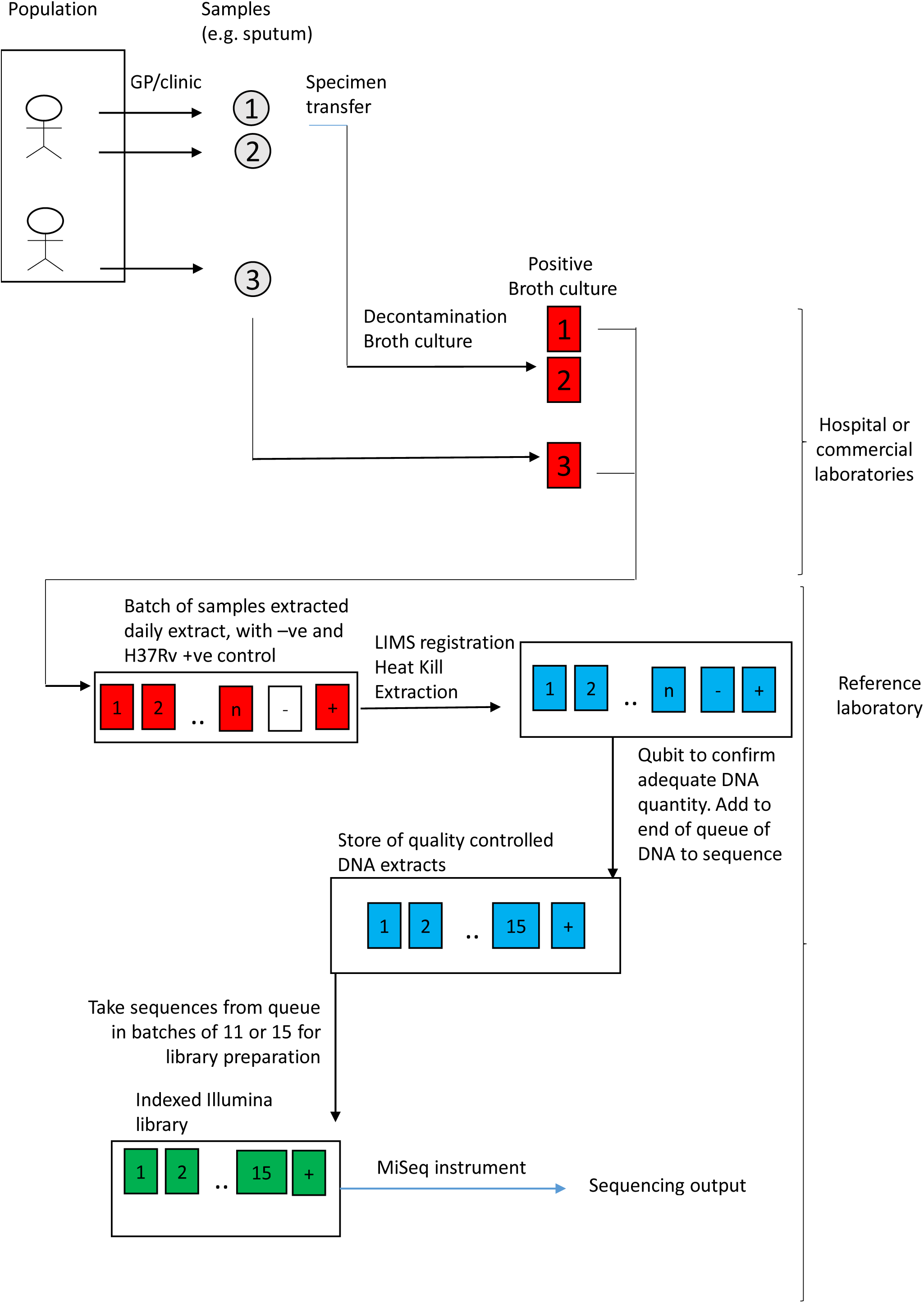
Laboratory and Bioinformatic processing

**Table 1.** Samples analysed

| Development Stage | Total sequences (samples and controls) | Date range | Number of individuals providing samples | MiSeq run range | Number of clinical samples | Number of MiSeq Runs |
| --- | --- | --- | --- | --- | --- | --- |
| Development | 938 | 20.04.2015-15.12.2015 | 630 | 101 .. 291 | 776 | 154 |
| Pre-Production | 1167 | 01.04.2016-06.12.2016 | 753 | 1152 ..1522 | 919 | 163 |
| Production | 2191 | 07.12.16-30.12.2017 | 1481 | 1523 .. 2307 | 1794 | 346 |

### Bioinformatic processing

The routine bioinformatics pipeline deployed by Public Health England has been previously described (15). Briefly, reads were first processed using the Mykrobe predictor tool, which identifies *Mycobacterium tuberculosis* using species specific k-mers (17). Specimens identified as containing *M. tuberculosis* were further processed (16), mapped to the H37Rv v2 genome (NC_000962.2) (18), as described (16), and vcf files generated using Samtools mPileup, with additional basecalling using GATK VariantAnnotator v2.1. A consensus base is called from high-quality bases provided one base accounts from >90% of the pileup; otherwise, the base is recorded as uncertain (‘N’) (15). In this analysis, high quality base counts (identified by the BaseCounts VCF tag) were extracted and summarised using code available at https://github.com/davidhwyllie/VCFMIX.

### Nucleotides identifying lineage

Coll *et al* (2) described the *M. tuberculosis* phylogeny and identified 62 sets of nucleotide positions defining the deep branches of the *Mycobacterium tuberculosis* lineage. At each position within a nucleotide set, in one particular clade, one nucleotide is uniquely present (i.e. is not present in any other of the known clades). These sets contain a median 108 nucleotide positions (range 1 – 898). In this analysis, we considered 55 branches, excluding branches 1.2, 3.1, 3.1.2, 4.1.2, 4.3.4.2.1, 4.6 and 4.7 because they contain fewer than 20 positions, making estimates of minor variation in these positions less reliable than estimates in other branches. We identified lineage using consensus basecalling in these 55 branches. If the signature SNV of a branch was called as uncertain, we called only to the level of the branch deeper (i.e. closer to the root) than the uncertain call. If more than one different lineage defining variant was called, or we could not call any lineage defining positions, we reported the samples as ‘lineage not defined’.

### Estimation of the minor variant frequency within a set

The minor variant frequency at a set of bases can be due to sequencing error, mapping error and/or *bona fide* inter-lineage mixtures (Supplementary Figure 1, panels A and B). Minor variant frequencies were determined as follows: if there are *n* bases in a lineage defining set, we count the high quality depths *d* at each base, e.g. if *n*=3 and *d*_1_ = 30, *d*_2_= 70, *d*_3_= 100, then the total depth 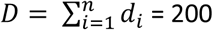. For each position, we also identify the most common base; the minor depth *m* is the total depth minus the most common base depth. If the minor depths are *m*_1_ = 3, *m*_2_= 7, *m*_3_= 10, then total minor depth 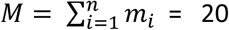; we estimate the minor allele fraction *p* in the set as M/D = 0.1.

### F2 and F47 metrics

If sequences from two different *M. tuberculosis* lineages are mixed together, then the sets which uniquely define these lineages will be mixed (depicted in Supplementary Figure 1C); there will be a minimum of two and maximum of eight sets affected (e.g. a Lineage 5/7 mixture will mix two sets of lineage defining nucleotides, a 2.1/4.2.1 will mix five sets, and a 4.1.1.1/3.1.2.1 mixture will mix 8 sets). Only if more than two samples are mixed will more than 8 sets be mixed. In this work, we describe two metrics reflecting mixing. Having computed the minor allele frequency estimates, *p*_1_, *p*_2_,.. *p*_55_ we can sort these in descending order, identifying the sets with the highest and lowest minor allele frequencies. We then estimate the minor variant frequency across the nucleotides in the top two (F2 metric) and lowest 47 sets (F47 metric) as described above. The underpinning assumptions are that mixtures of biological origin are most likely to occur between two lineages, and therefore F2 is the most sensitive metric for identifying these. Since between two and 8 sets will be mixed in such genuine co-infections, then the lowest 47 (55 - 8) sets will not be mixed, and thus the F47 metric is more sensitive for identifying laboratory contamination involving more than two samples.

### Regression modelling

Because of high leverage by a small number of observations, we used quantile regression to estimate the relationship between median values of log-transformed non-callable base numbers and log-transformed F47, using the quantreg R package (R 3.3.1).

### Ethical framework

Only anonymised data was used in this work; ethical approval is not required.

### Data availability

The data analysed is available at https://ora.ox.ac.uk/objects/uuid:5e4ec1f8-e212-47db-8910-161a303a0757.

## RESULTS

### Samples studied

4,156 samples were included since they (i) were identified using MyKrobe (17) as belonging to the *M. tuberculosis* complex, and (ii) had at least 0.5 × 10^6^ read pairs mapped to the H37Rv reference genome, a criterion reflecting successful DNA extraction and sequencing. These sequences were highly diverse, originating from six branches of lineage 1 (n=320), five branches from lineage 2 (n=278), five branches of lineage 3 (n=1,010), thirty lineage 4 branches (n=2,266). 106 samples were from *M. bovis* or *africanum*, and 176 did not have their lineages defined.

The laboratory processes operated under three different phases: in the first, *development*, laboratory processes were being actively refined; in the second, *pre-production*, laboratory processes were fixed and controlled by standard operating procedures, with version controlled changes. The third *production* state was similar to the second, except that the process had received ISO15189 accreditation.

### Variation in lineage defining positions in H37Rv controls

The F2 mixture metric reflects the estimated mixture in the two most mixed lineage defining sets; in the H37Rv controls, this follows a distribution skewed to the right (Figure 2A). The MiSeq runs with H37Rv controls with F2 mixture metrics in the top 5% (Fig. 2A) are temporally clustered (red lines on Fig. 2B, C) with a number of examples in the Development phase. Among clinical (non-control) samples, variation in F2 metric is explained in part by MiSeq run (Kruskal-Wallis test, p < 10^−16^), and a strong correlation exists between the F2 metric in H37Rv controls and that in clinical samples on the same plate (ρ = 0.61, 95% CI 0.56-0.61, Spearman’s Rank Correlation). That is, in plates with elevated F2 metrics in the H37Rv control, the clinical samples are more likely to have elevated F2, as is evident visually (e.g. Fig 2B and C, around run 2301).

**Figure 2.**
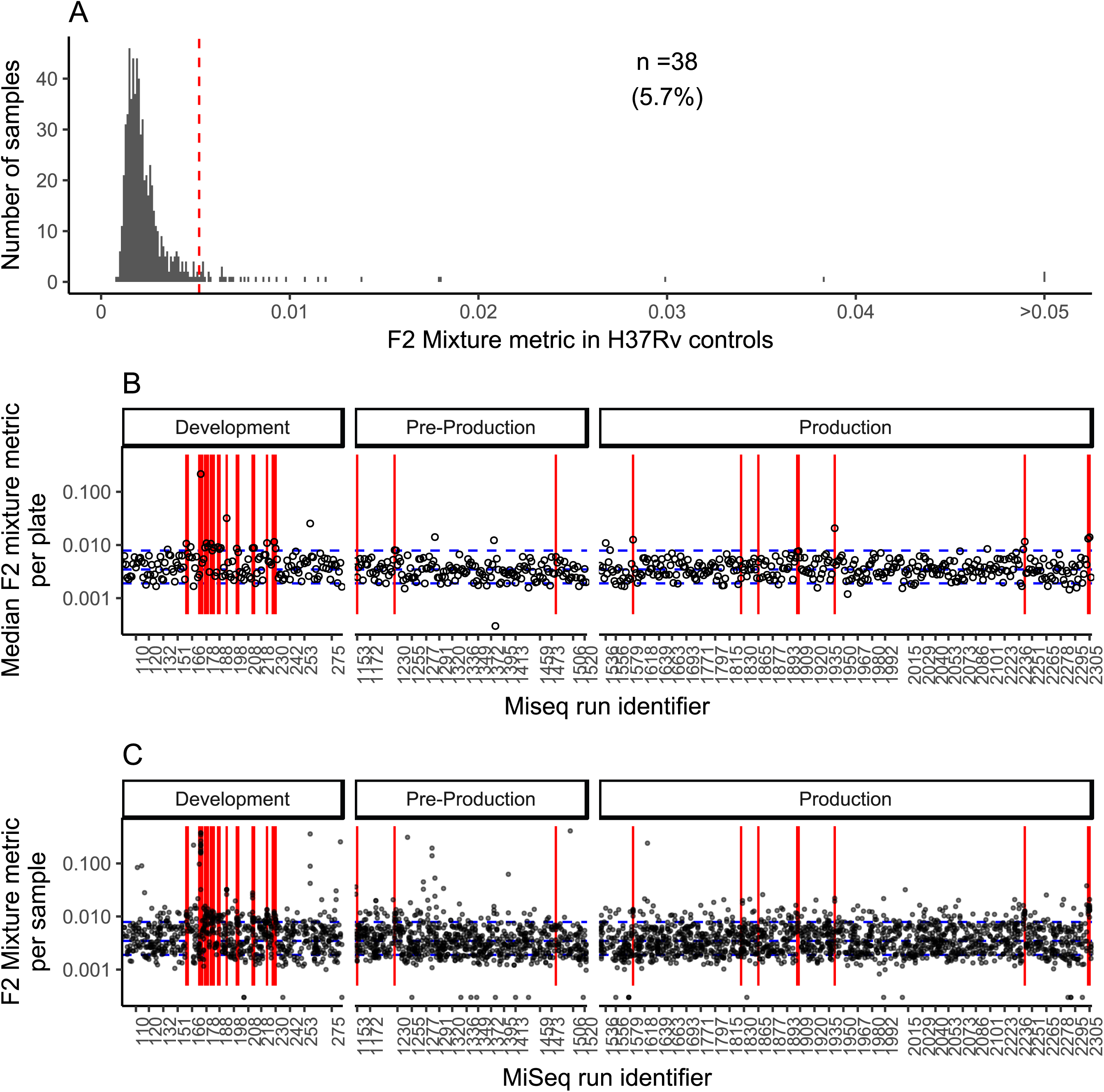
Mixtures in H37Rv controls. Histogram showing the F2 metric, which reflects the mixture in the two most mixed lineage associated sets, in H37Rv control DNA (A). Median F2 metric among clinical samples other than H37Rv is shown in (B); red lines indicate that the F2 mixture metric in H37Rv controls is raised (as shown in A). (C) shows F2 metric for each *M. tuberculosis* sequence from a clinical sample.

### Different patterns of mixtures were observed during Development

We ordered specimens first by the order of the plates analysed and the order in which the bioinformatics processing was completed, which is the order that an automated quality control monitoring system would encounter output. During the Development phase (Fig. 3), blocks of samples derived from runs with elevated mixtures in the H37Rv control are seen (red bars in Fig. 3A), coincident with clear increases in both F2 and F47 metrics (Fig. 3B, C), reflecting elevations in mixed bases across most or all lineage defining positions (Fig. 3D). These blocks of samples typically span multiple MiSeq runs (Fig 3A,B). In addition to the blocks of samples with elevated F2 and F47 metrics, we also observed small numbers of single samples with elevated F2, but not F47, metrics (Fig. 3B, Arrow). This latter pattern is expected in cases of inter-lineage mixtures of only two samples (Supp. Fig 1, and Methods). These patterns were also seen in the subsequent phases (Supplementary Figures S2–5, yellow dots).

**Figure 3.**
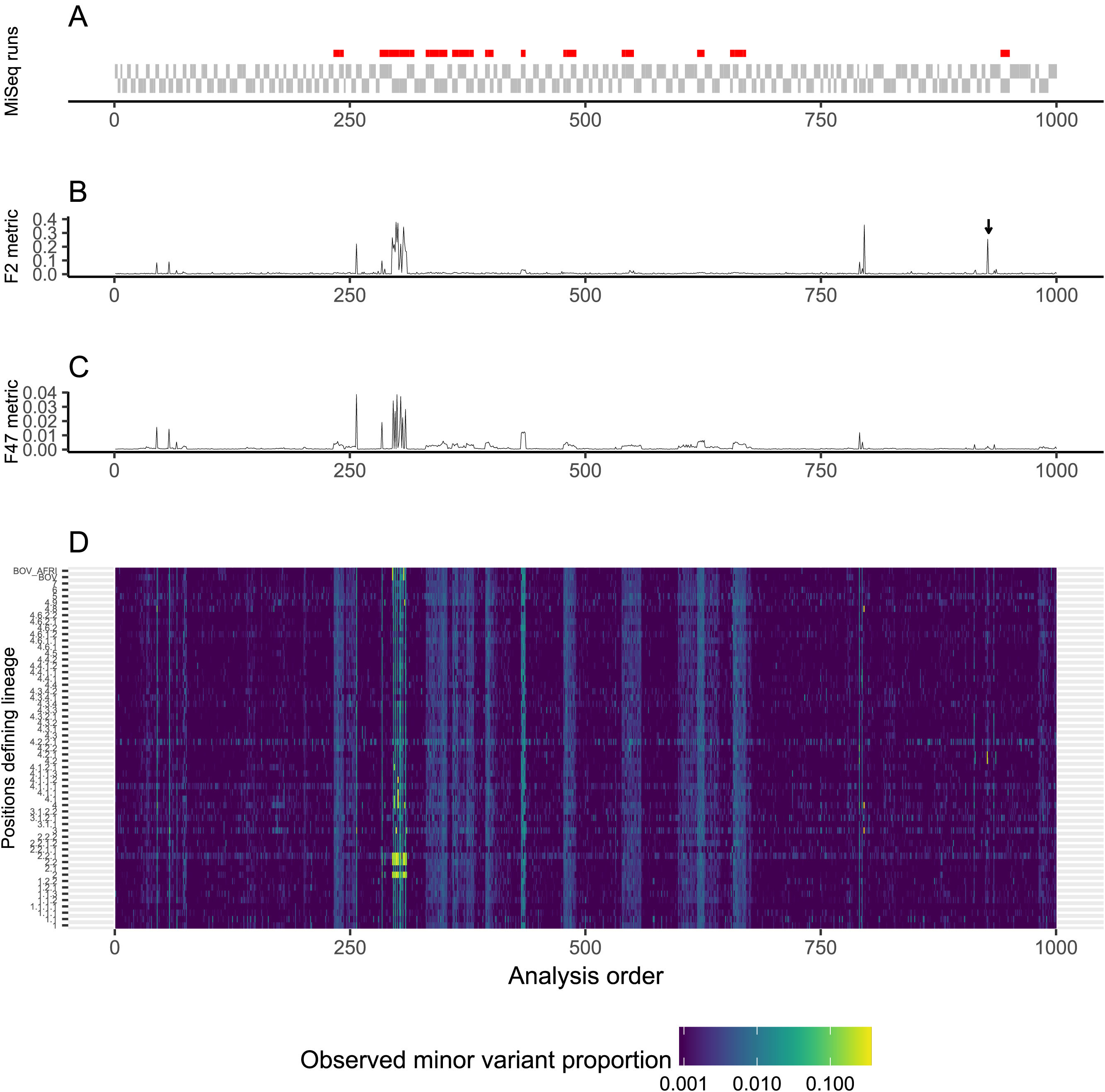
Mixture metrics in the Development phase. Samples arranged first by the order of the MiSeq runs (depicted as solid gray blocks, in A), and the order bioinformatics processing completed. Only samples yielding *M. tuberculosis* are shown, which is why some blocks in A are longer than others. If the H37Rv control samples had increased F2 statistics, a red bar is shown above each sample in A. We depicted the F2 (B) and F47 metrics (C), as well as the estimated mixture F in each of the 58 lineage defining sets (D). The arrow illustrates a sample with elevated F2, but low F47 metric.

### Mixtures of multiple lineages are common

Based on the pattern observed in the Development phase, we categorised samples as having one of (i) neither F2 nor F47 raised, (ii) raised F47, or (iii) raised F2 without raised F47. We defined a raised F2 and F47 as more than 10x and 5x the respective median metric during Development in all control and clinical samples, cutoffs which correspond to 4.7% (F2) and 0.2% (F47) minor variant frequencies across the relevant lineage-defining sets, respectively. In the Pre-production and Production phases, F2 and F47 values below these thresholds (reflecting unmixed samples) were observed in 97.5% of the samples studied, raised F47 and F2 values (reflecting a mixture of multiple samples) in 2.5%, while six samples (0.001%) had raised F2 but normal F47 values (Table 2).

**Table 2.** Detection of mixtures in clinical samples

| Development Stage | Neither F2 nor F47 raised | F2 raised, but F47 normal | F47 raised |
| --- | --- | --- | --- |
| Pre-Production (n=919) | 900 (98%) | 5 (0.003%) | 14 (1.1%) |
| Production (n=1794) | 1741 (97%) | 1 (0.001%) | 52 (2.8%) |

### Isolated F2 metric elevation is rare

Isolated elevation of F2 is expected if bacteria from two different lineages are mixed. In Fig. 4, we show the minor variant frequencies from all samples from the six individuals with raised F2, but normal F47, metrics. In one case, patient 3, two technical repeats of the same sample (#2) showed the same pattern, as did a separate sample taken contemporaneously. In other cases (patients 1, 5, 6) the mixed pattern was only observed in one out of two positive samples taken on the same day, and in two cases (2, 3) only a single sample was positive. Thus, between 1 and 6 samples of the 4,156 studied may truly reflect mixed co-infections.

**Figure 4.**
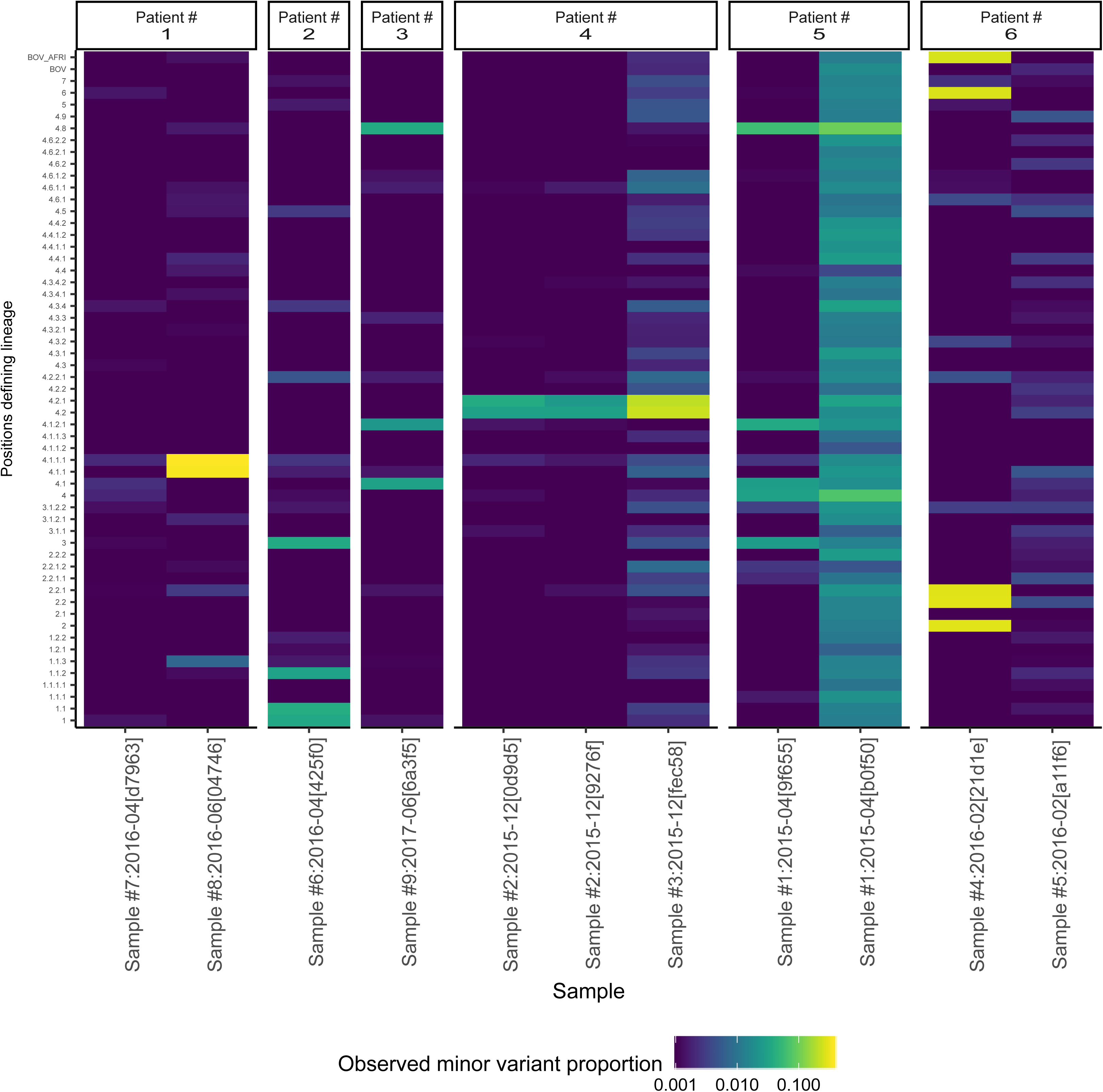
Consistency of isolated F2 elevation in individuals. Six individuals with elevated F2, but not F47, statistics were identified during the pre-production and production phases. The observed minor variant proportion for all deep branches analysed are shown in a heatmap. For example, patient #4 had two samples taken in December 2015; sample 2 was analysed twice (sequencing ids 0d9d5,9276f) and sample 3 once. A similar pattern of minor variation is seen in all three samples.

### Impact of inter-lineage variation on basecalling

One obvious question is whether very low level cross-contamination impacts the consensus sequence which can discerned from the pileup. As cross-contamination increases, at some point minor variant frequencies in some parts of the genome will start to rise above the 10% cutoff specified by the basecalling algorithm. The numbers of uncalled bases will then rise; this relationship can be observed in Figure 5: the number of uncallable bases rises rapidly when F47 exceeds the cutoff value, but only slowly below it. Below the cutoff value of 4.7% (red line in Fig. 5), which is 10 times the median (black line in Fig. 5), the number of uncallable bases increased by 1.25 fold (95% CI 1.22, 1.28) for every 10-fold increase in F47; above the cutoff, the corresponding increase was 9.24 (95% CI 5.5, 11.2).

**Figure 5.**
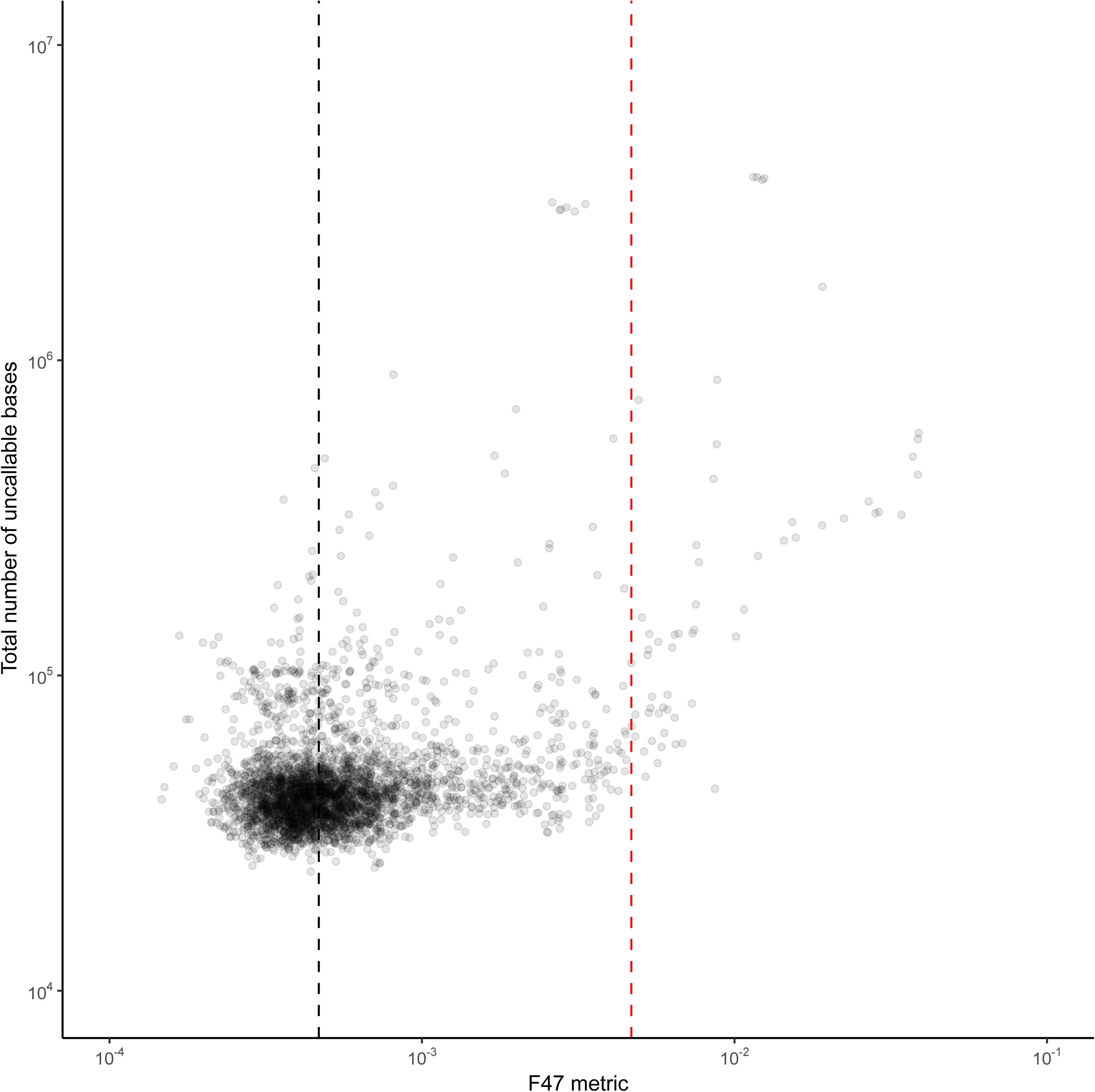
F47 metric and the number of callable bases. The relationship between the F47 metric and the number of uncallable bases. The red line corresponds to the cutoff used to define F47 as being elevated.

## DISCUSSION

In this work, we describe methods for monitoring the presence of mixtures of *Mycobacterium tuberculosis* of different lineages. It assumes that multiple lineages and sublineages of *M. tuberculosis* are being sequenced contemporaneously; this is the case in our setting, and is also true globally (19, 20). Using single nucleotide variants, each of which uniquely defines a branch in the phylogenetic tree of *M. tuberculosis*, we can show two patterns of mixtures. The first, which occurred in about 2.5% of samples during the pre-production and production phases of our project, is indicative of multiple samples being mixed together, since mixtures are seen in most or all of the lineage defining branches. This occurred in batches, was characterised by cross-contamination at levels of less than 1%, and can be monitored by a metric we term F47. This pattern likely reflects process failures. The strength of the F47 metric is that the depth analysed is very high, as about 5,000 nucleotides typically contribute across the lineage-defining sets included in it. If there is a sequencing depth of 50-100 at each of these, the effective sequencing depth analysed is of the order of 25,000 – 50,000, making detection of minor variation at sub- 1% levels readily feasible with high statistical confidence.

Such low level cross-contamination, as observed during our production process and illustrated in Supplementary Figures 2–4, is likely to have minimal influence on inference drawn from the sequence, unless highly sensitive assays for heteroresistance are required. A sensitive metric such as F47 will, however, allow early detection of emerging problems and allow review of process, as part of continuous quality improvement.

A second class of mixture, which we found was rarely detected in this setting, is compatible with co-infection with two organisms of differing lineage within the patient. This kind of mixture is clinically relevant (7, 11), and may be under-detected using the laboratory process we describe here, since culture based amplification can reduce diversity in the sample inoculated (10). Its frequency may rise if direct-from-sample sequencing is employed, or if samples from high-endemicity areas are studied, but here we identified only one probable case of such mixtures, and five other possible cases. Confirmatory approaches are available: microbiological techniques conducted separately on multiple picks from the same samples have been used as validation(3); a limitation of this study is that we could not undertake such work as only multiply sub-cultured stored isolates exist for historical samples. Techniques for reconstructing the contributing sequences also have been described in detail (21–23), and we did not study them here. Another limitation is that we were not able to consider six of the lineage defining sets in Coll *et al* because they covered <20 nucleotide positions and therefore we considered that they did not contain sufficient information to be used in F2 or F47 metrics. A consequence is that our method would not identify mixtures of samples if they only involved mixtures in these excluded branches.

The clinical of use of bacterial genome sequencing is rising (16, 17, 24), and given the importance of *M. tuberculosis* and the complexity of treatment, *M. tuberculosis* has been one of the first organisms tackled (15). The processes followed involve multiple steps at which the opportunity for cross-contamination exists. The availability of tools monitoring critical aspects of laboratory process is required for accreditation under ISO15189, and the F2 and F47 metrics described here will contribute to this.

## ACKNOWLEDGEMENTS

This study is supported by the Health Innovation Challenge Fund (a parallel funding partnership between the Wellcome Trust [WT098615/Z/12/Z] and the Department of Health [grant HICF-T5-358]) and NIHR Oxford Biomedical Research Centre. Professors Derrick Crook, Tim Peto and Sarah Walker are affiliated to the National Institute for Health Research Health Protection Research Unit (NIHR HPRU) in Healthcare Associated Infections and Antimicrobial Resistance at University of Oxford in partnership with Public Health England. Professor Crook is based at University of Oxford. Professor Tim Peto is an NIHR Senior Investigator. The views expressed are those of the author(s) and not necessarily those of the NHS, the NIHR, the Department of Health or Public Health England. The sponsors of the study had no role in study design, data collection, data analysis, data interpretation, or writing of the report. The corresponding author had full access to all the data in the study and had final responsibility for the decision to submit for publication.

## Supplementary Figures

**Supplementary Figure 1.**
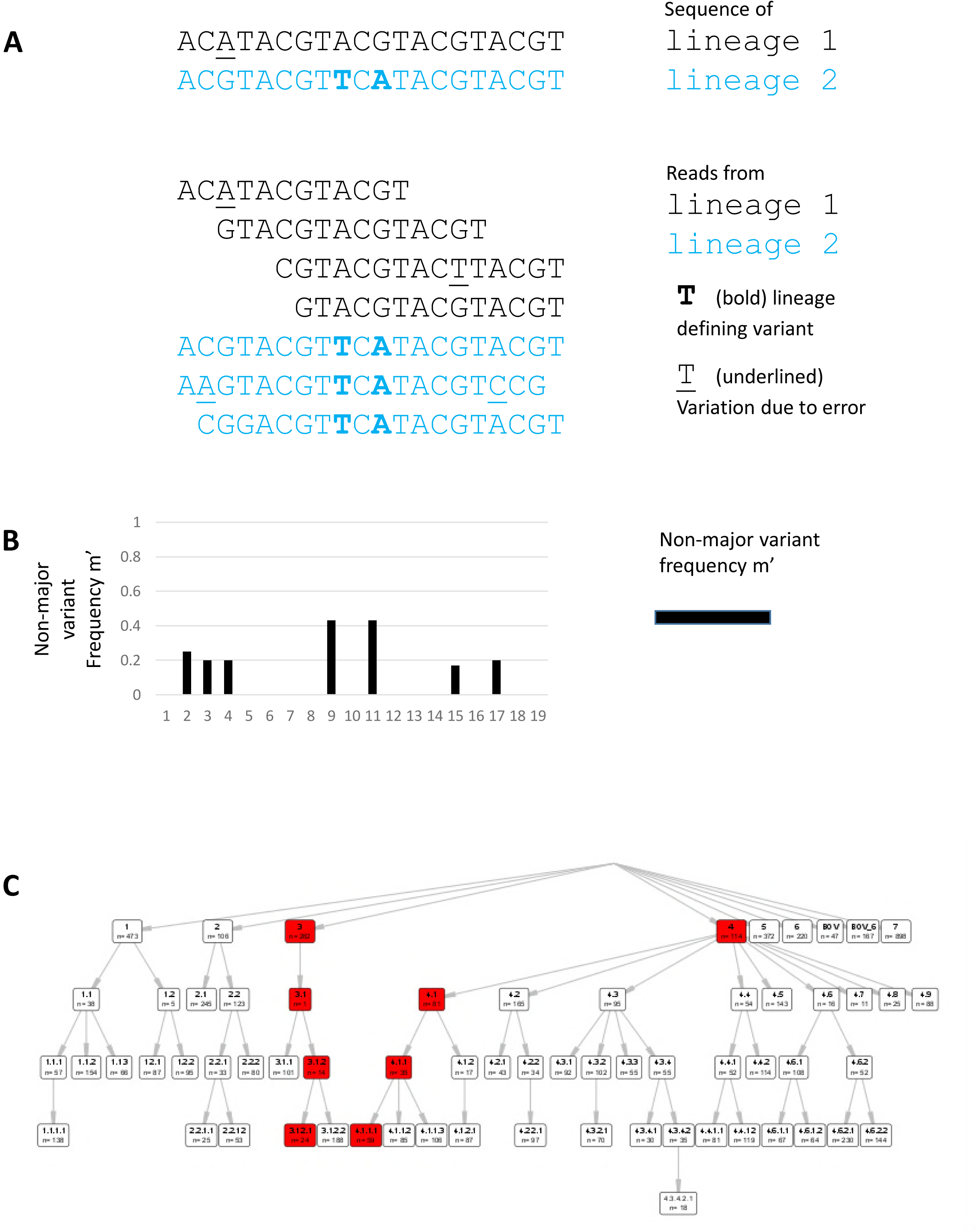
Illustrates the computation of variation between samples of two different lineages, Lineage 1 (black text) and Lineage 2 (blue text) (A). When a mixture of these samples is present, and mapped to a reference sequence, a major base and minor base(s) are present in the pileup (B). Variation may be due to either sequencing error (underlined) or to lineage associated variation; the non-major variant frequency included both classes of variation. Lineage defining sites, as defined by Coll *et al* (2), mark branches of the phylogenetic tree. If a lineage 4.1.1.1 *M. tuberculosis* is mixed with a lineage 3.1.2.1 *M. tuberculosis*, eight sets of lineage defining sites will be mixed (red boxes).

**Supplementary Figures 2-5.**
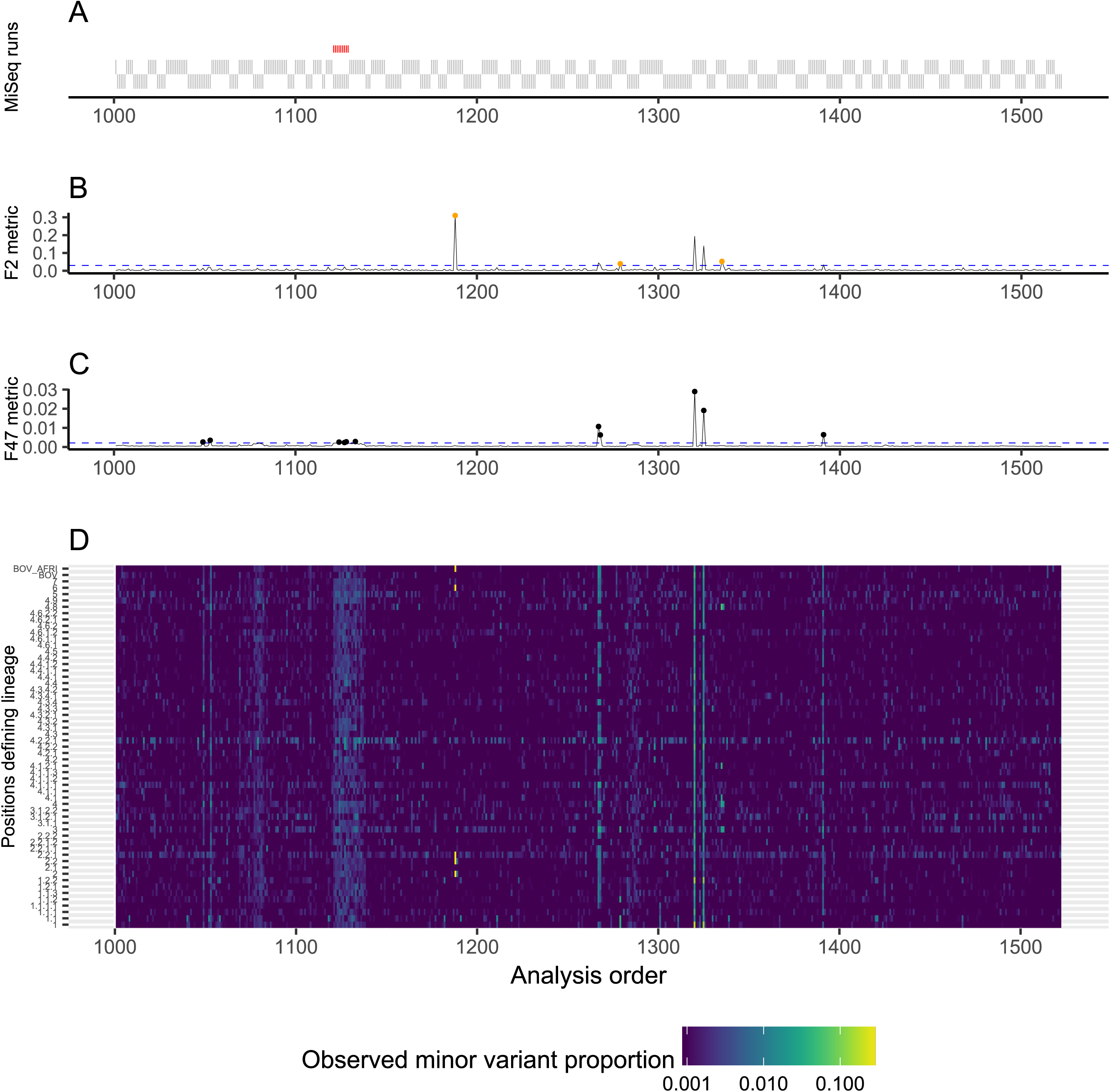
These illustrate mixture patterns observed during the Production stage. The layout is similar to Figure 3; samples arranged first by the order of the MiSeq runs (depicted as solid gray blocks, in A), and the order bioinformatics processing completed. Only samples yielding *M. tuberculosis* samples are shown, which is why some blocks in A are longer than others. If the H37Rv control samples had increase F2 statistics, a red bar is shown above each sample in A. We depicted the F2 metric (B) and F47 metrics (C), as well as the estimated mixture F in each of the 58 lineage defining sets (D).

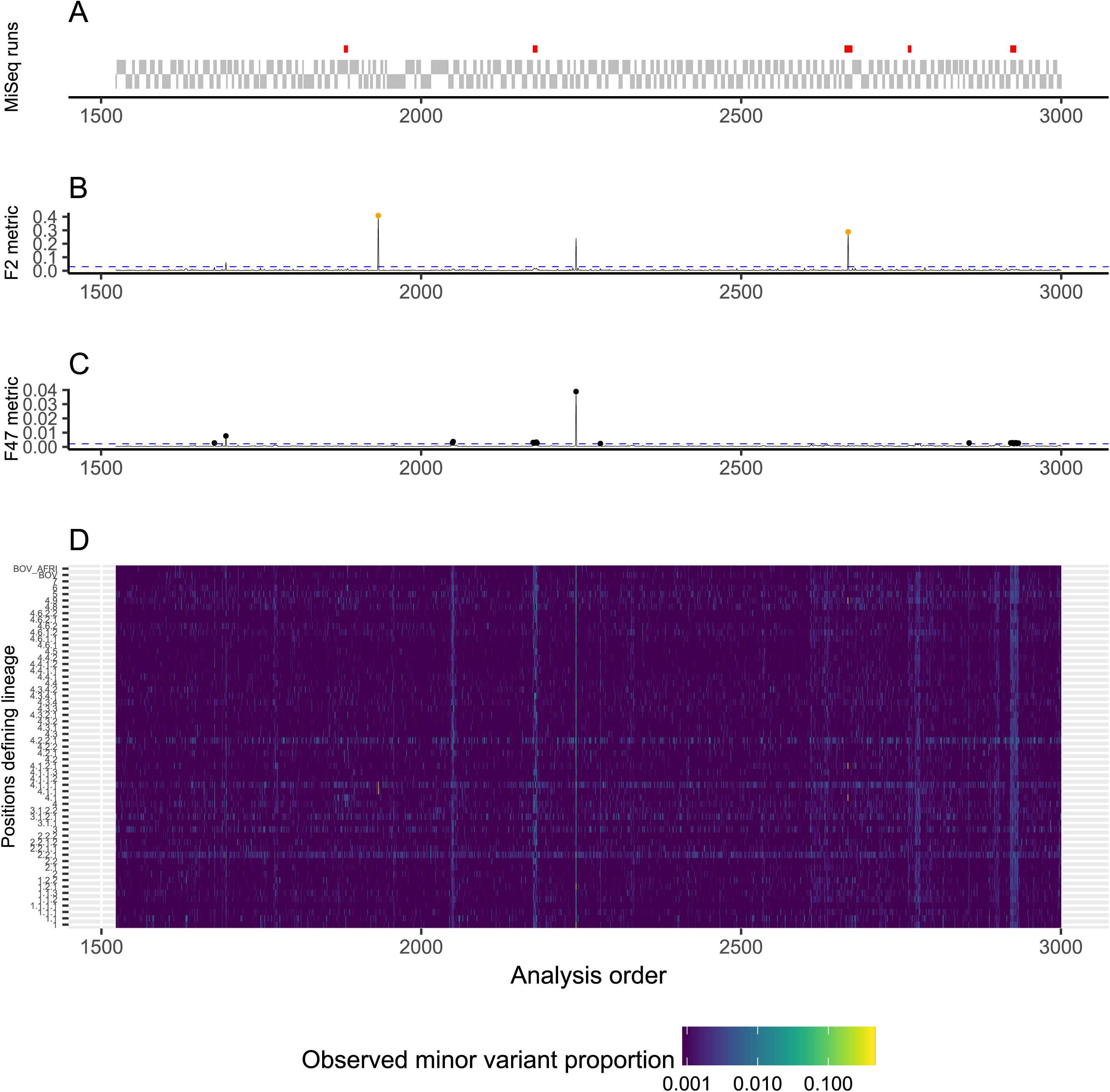

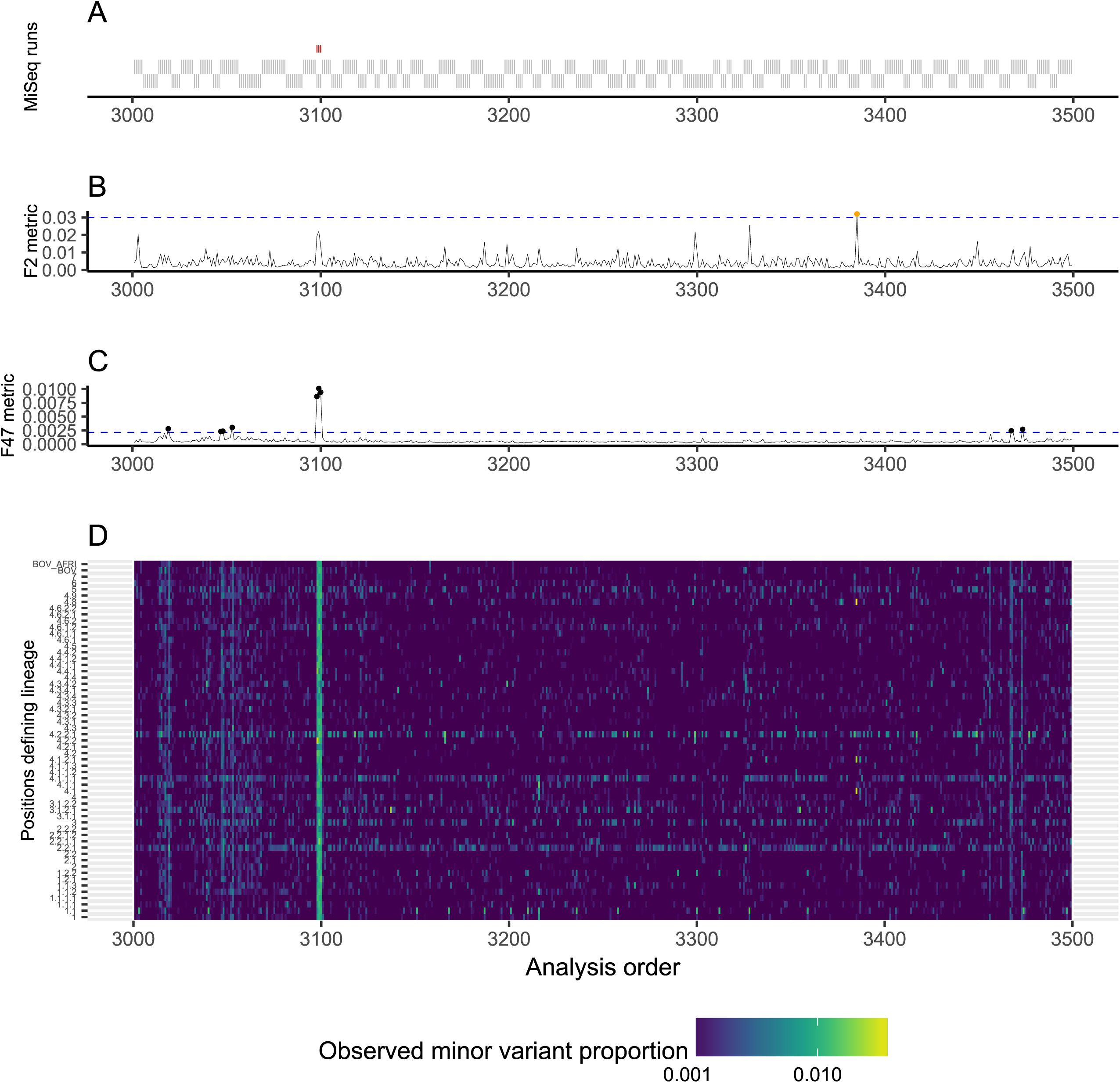

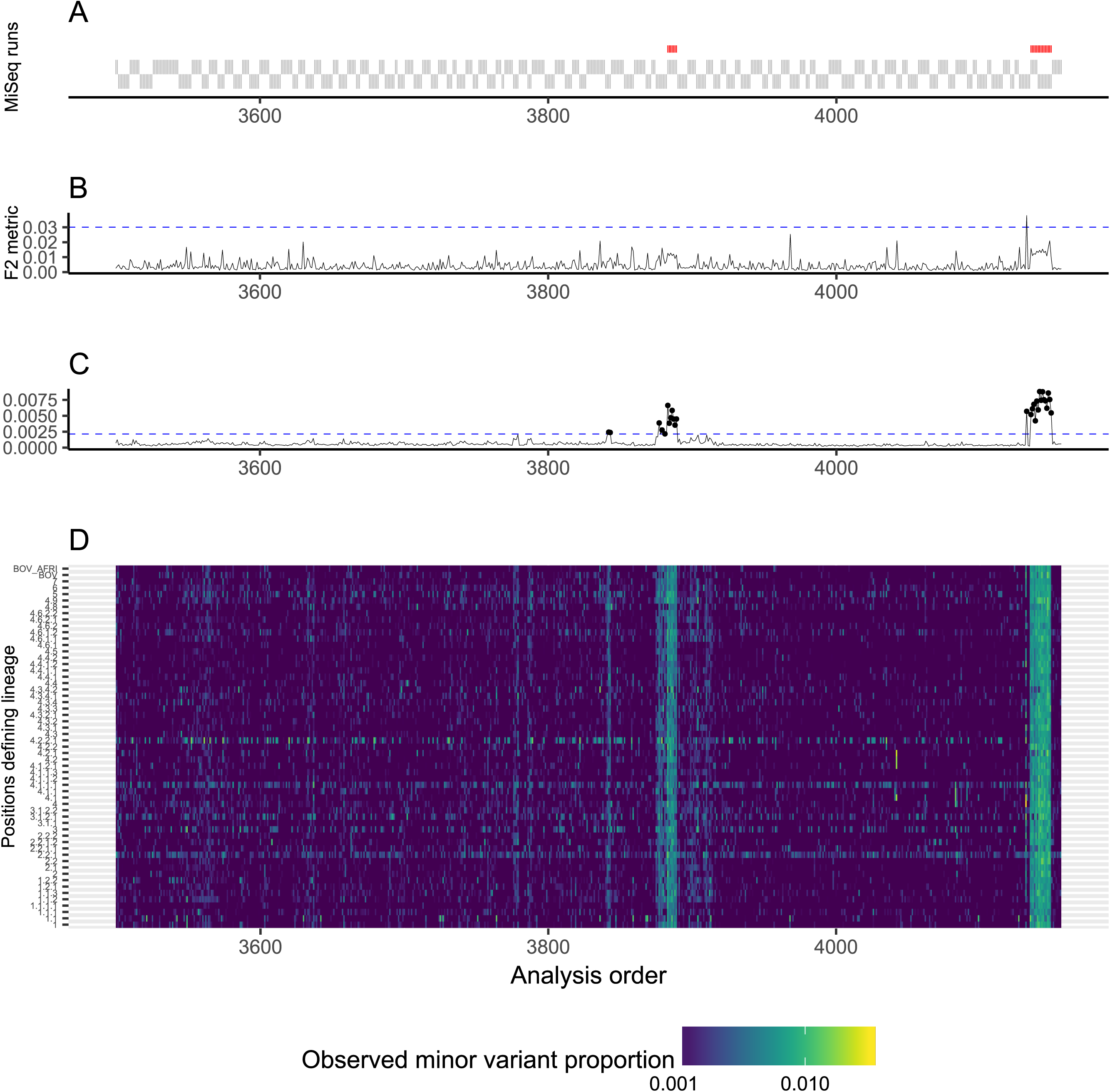

